# Knockout of Dectin-1 does not modify disease onset or progression in a MATR3 S85C knock-in mouse model of ALS

**DOI:** 10.1101/2024.04.18.590117

**Authors:** Justin You, Katarina Maksimovic, Karin Chen, Jooyun Lee, Anneka Schoeppe, Jhune Rizsan Santos, Mohieldin M. M. Youssef, Michael W. Salter, Jeehye Park

## Abstract

Microglia have been increasingly implicated in neurodegenerative diseases, including amyotrophic lateral sclerosis (ALS). Dectin-1, encoded by the *Clec7a* gene, is highly upregulated in a specific microglial response state called disease-associated microglia (DAM) in various neurodegenerative conditions. However, the role of Dectin-1 in ALS is undetermined. Here, we show that *Clec7a* mRNA upregulation occurs in central nervous system (CNS) regions that exhibit neurodegeneration in a MATR3 S85C knock-in mouse model (*Matr3^S85C/S85C^*) of ALS. Furthermore, a significant increase in the number of Dectin-1^+^ microglia coincides with the onset of motor deficits, and this number increases with disease severity. We demonstrate that the knockout of Dectin-1 does not affect survival, motor function, neurodegeneration, or microglial responses in *Matr3^S85C/S85C^*mice. These findings suggest that Dectin-1 does not play a role in modifying ALS onset or progression but could potentially serve as a valuable biomarker for ALS severity.

**Subject areas:** Physiology; Molecular biology; Neuroscience; Immunology

**Highlights:**

- *Clec7a* upregulation is confined to central nervous system regions that exhibit overt neurodegeneration in a MATR3 S85C knock-in mouse model of ALS
- The appearance of Dectin-1^+^ microglia coincides with the onset of motor deficits, and its number increases with disease progression
- Knockout of Dectin-1 does not modify survival, motor deficits, neurodegeneration, or microglial responses in MATR3 S85C knock-in mice

## Introduction

Microglia are highly plastic and can exist as several states with unique functions in health and disease^1–4^. In neurodegenerative diseases, a specific microglial state termed disease-associated microglia (DAM) appear in the presence of neurodegenerative cues, including apoptotic neurons, myelin debris, and toxic proteins such as amyloid beta (Aβ) and pathological TDP-43^5–8^. DAM display a specific molecular signature with upregulated expression of genes involved in phagocytic, adhesion, lysosomal, and lipid metabolism pathways, including *Clec7a*, *Itgax*, *Lpl*, *ApoE* and *Trem2*^3,5^. In particular, *Trem2* is essential in the transition of homeostatic microglia to a DAM or DAM-like state in the presence of these neurodegenerative cues^5,6,9,10^. *Trem2* knockout blocks microglial transition to DAM and has largely been shown to hinder microglial phagocytic function and worsen neurodegenerative disease phenotypes in mouse models of Alzheimer’s disease (AD), multiple sclerosis (MS) and amyotrophic lateral sclerosis (ALS)^7–9,11,12^. Furthermore, overexpression of *Trem2* ameliorates the disease phenotypes in an AD mouse model^13^. These findings suggest that DAM may play a neuroprotective role and represent a promising therapeutic target in neurodegenerative diseases. However, the function of other DAM genes, such as the *Clec7a*, remains elusive.

*Clec7a*, which encodes for Dectin-1, is among the most characterized C-type lectin receptor in myeloid cells, including macrophages, monocytes, neutrophils, and dendritic cells, and was initially identified to play important roles in the innate immune response^14,15^. Recently, there has been an increasing appreciation of the roles of Dectin-1 in microglia^16^. Results from single-cell RNA-sequencing revealed that *Clec7a* is not expressed in homeostatic microglia, but rather in several other microglial states, including DAM^5^ and white matter-associated microglia (WAM) during aging^10^.

Several studies have investigated the role of Dectin-1 in various neurological disorders. In MS mice, knockout of Dectin-1 exacerbated disease severity, whereas an injection of a Dectin-1 agonist ameliorated disease severity^17^. In AD mice, intrahippocampal injections of a Dectin-1 agonist decreased Αβ load^18^. Another paper supported these results, showing that intraperitoneal injection of anti-Dectin-1 antibody in AD mice partially rescues microglial response to Aβ^9^. In addition, intraocular injection of a Dectin-1 agonist induced axonal regeneration after optic nerve crush injury^19–21^. Although these studies suggest a beneficial role of Dectin-1 signaling in various neurodegenerative diseases, there have also been several conflicting studies. A recent study showed that knockout of Dectin-1 ameliorated cognitive deficits and neurodegeneration in AD mice^22^. Similarly, adeno-associated virus-mediated knockdown of Dectin-1 alleviated motor deficits and neurodegeneration in a Parkinson’s disease (PD) rat model^23^. Furthermore, pharmacological inhibition of Dectin-1 partially rescued the disease phenotypes in mouse models of ischemic stroke^24^ and intracerebral hemorrhage^25^. Together, these findings suggest that Dectin-1 could also play a detrimental role in neurological disorders. These conflicting results highlight the need for further research to understand the disease-modifying roles of Dectin-1. In particular, the role of Dectin-1 in ALS has not yet been investigated.

ALS is a devastating neurodegenerative disease characterized by the progressive loss of motor function. Patients typically survive for 2-5 years after disease onset, and there is currently no cure. Mutations in *MATR3* were first identified in 2014 to cause ALS, including the familial S85C mutation, the most prevalent mutation in *MATR3*^26^. MATR3 is a DNA- and RNA-binding protein involved in transcription and RNA metabolism^27,28^. To investigate the impact of the S85C mutation in *MATR3*, we have previously generated a MATR3 S85C knock-in mouse line (*Matr3^S85C/S85C^*)^29^. We have previously demonstrated that *Matr3^S85C/S85C^* mice recapitulate ALS-like features, including a shortened lifespan, motor deficits, loss of Purkinje cells in the cerebellum, degeneration of lower motor neurons in the spinal cord, and microglial responses^29,30^. We have shown from RNA-sequencing data that *Clec7a* was the topmost upregulated gene in the cerebellum and lumbar spinal cord of these mutant mice^29^. Therefore, we sought to utilize this model to investigate the role of Dectin-1 in ALS. In this study, we show that *Clec7a* is upregulated in the affected regions such as cerebellum and spinal cord but not in other central nervous system (CNS) regions or in peripheral tissues. We found that the appearance of Dectin-1^+^ microglia coincides with the onset of motor deficits and that the number of Dectin-1^+^ microglia increase with disease severity. To investigate the role of Dectin-1 in ALS, we generated *Matr3^S85C/S85C^* mice lacking Dectin-1. We demonstrate that the knockout of Dectin-1 does not modify survival, motor function, neurodegeneration, or microglial responses in *Matr3^S85C/S85C^* mice, suggesting that Dectin-1 does not affect ALS onset or progression.

## Results

### *Clec7a* is upregulated in the affected CNS regions, such as cerebellum and spinal cord of *Matr3^S85C/S85C^* mice

The transition of homeostatic microglia to DAM and the consequent upregulation of *Clec7a* have been shown to occur in response to neurodegenerative cues^3^. Our previous RNA sequencing results showed that *Clec7a* is upregulated in the cerebellum and lumbar spinal cord of *Matr3^S85C/S85C^* mice^29^. Both the cerebellum and lumbar spinal cord are the affected regions in *Matr3^S85C/S85C^*mice as they exhibit degenerative phenotypes in Purkinje cells and lower motor neurons, respectively^29^. For further validation of *Clec7a* differential expression, we extracted RNA from the cerebellum and spinal cord of *Matr3^+/+^* and *Matr3^S85C/S85C^*mice at 30 weeks of age and performed RT-PCR. Indeed, there was a significant upregulation of *Clec7a* in the cerebellum and spinal cord of *Matr3^S85C/S85C^* mice compared to *Matr3^+/+^* mice (Figures 1A and 1B). Next, we assessed whether *Clec7a* was upregulated in other brain regions as well as in the peripheral tissues of the mutant mice. We found no significant difference in *Clec7a* expression in other areas of the brain, spleen, or tibialis anterior muscle between *Matr3^S85C/S85C^*and *Matr3^+/+^* mice (Figures 1A and 1B). These findings demonstrate that the upregulation of *Clec7a* is confined to the affected CNS regions in *Matr3^S85C/S85C^* mice.

**Figure 1.**
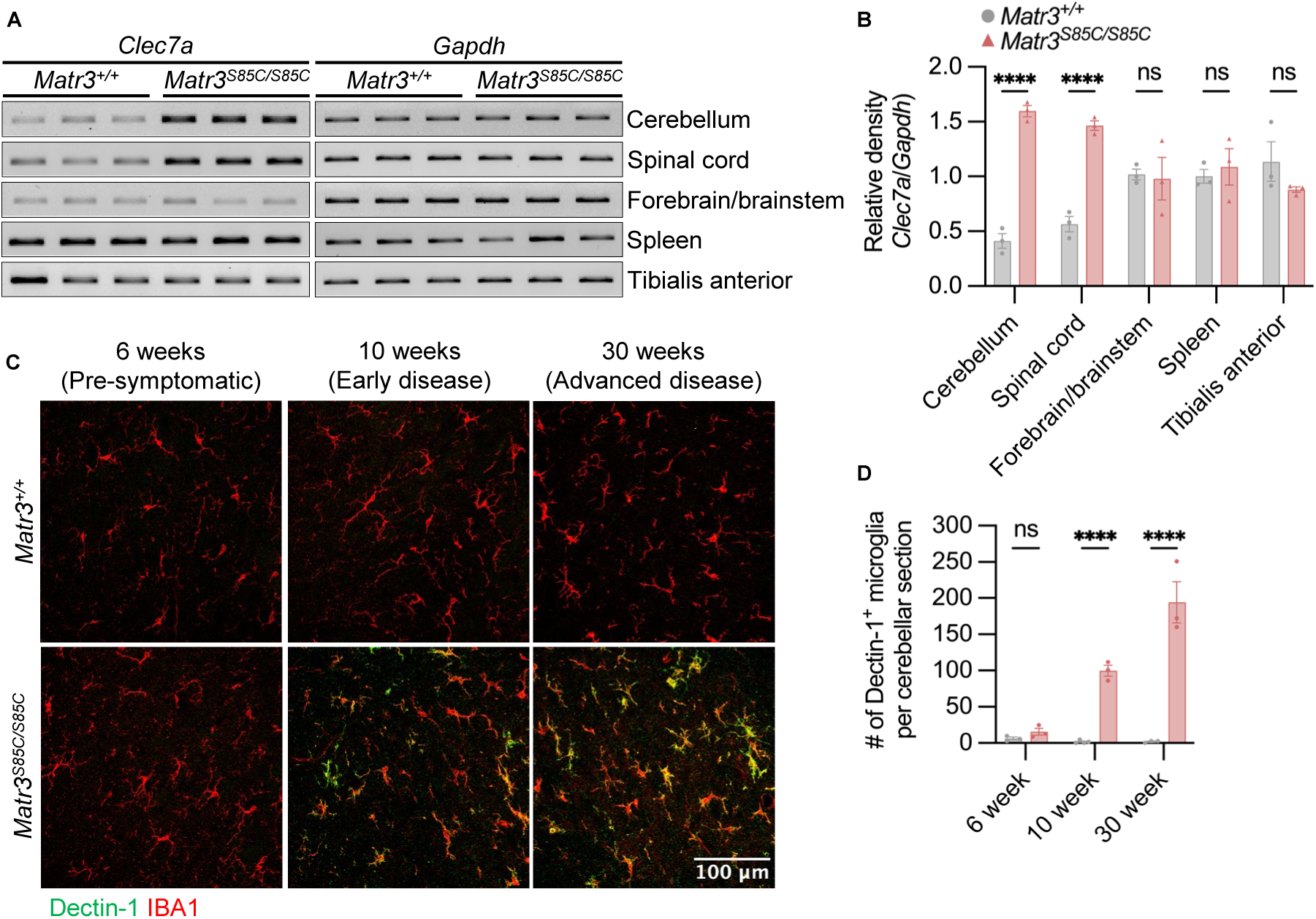
*Clec7a* upregulation is confined to the cerebellum and spinal cord in *Matr3^S85C/S85C^* mice. (A) Representative images and (B) quantification of RT-PCR results showing the mRNA expression levels of *Clec7a* and *Gapdh* from various tissues obtained at 30 weeks of age (*Matr3^+/+^*, *n* = 3; *Matr3^S85C/S85C^*, *n* = 3). *Clec7a* expression is normalized to *Gapdh* expression. Bar heights depict mean ± SEM, with each dot representing a single animal. Significance was determined using an ordinary two-way ANOVA corrected for multiple comparisons using the Šídák method. (C) Representative images of the deep cerebellar nuclei showing Dectin-1 (green) and IBA1 (red) staining for *Matr3^+/+^*and *Matr3^S85C/S85C^* mice at the indicated ages. (D) Quantification of the total number of Dectin-1^+^ microglia per cerebellar section (*Matr3^+/+^*, *n* = 3; *Matr3^S85C/S85C^*, *n* = 3). Bar heights depict mean ± SEM, with each dot representing a single animal. Significance was determined using an ordinary two-way ANOVA corrected for multiple comparisons using the Šídák method.

### Increased number of Dectin-1^+^ microglia coincides with disease onset and correlates with disease progression in *Matr3^S85C/S85C^* mice

To assess when Dectin-1^+^ microglia start to appear in *Matr3^S85C/S85C^*mice during disease progression, we conducted immunostaining on cerebellar sections with Dectin-1 and IBA1 antibodies at various disease stages, including the pre-symptomatic stage (6 weeks), early disease stage (10 weeks), and advanced disease stage (30 weeks). Notably, we previously defined the early disease stage as 10 weeks of age, corresponding to when *Matr3^S85C/S85C^* mice start to exhibit motor deficits compared to *Matr3^+/+^*mice^29^. We found that the number of Dectin-1^+^ microglia is significantly increased in *Matr3^S85C/S85C^*compared to *Matr3^+/+^* mice at the early and advanced disease stages but not at the pre-symptomatic stage (Figures 1C and 1D). Furthermore, the number of Dectin-1^+^ microglia further increased in the advanced disease stage compared to the early disease stage, correlating with disease severity (Figure 1D). Notably, this significant increase in Dectin-1^+^ microglia was present throughout the cerebellum in the mutant mice (Figure S1). However, we found no appreciable numbers of Dectin-1^+^ microglia in the cerebellum of *Matr3^+/+^* mice at 6, 10, or 30 weeks of age (Figures 1C and 1D). These findings suggest that the appearance of Dectin-1^+^ microglia coincides with the onset of motor deficits and increase with disease progression in ALS.

### Knockout of Dectin-1 does not impact motor phenotypes or survival of *Matr3^S85C/S85C^* mice

Next, we sought to investigate the impact of Dectin-1 knockout on ALS onset and progression. Dectin-1 knockout mice (*Clec7a^-/-^* mice) were viable and fertile^31^, and we confirmed the loss of *Clec7a* mRNA and protein expression in *Clec7a^-/-^* mice by RT-PCR and immunostaining, respectively (Figure S2). We then crossed MATR3 S85C KI mice to Dectin-1 knockout mice (*Clec7a^-/-^*)^31^. We found that *Matr3^S85C/S85C^ Clec7a^-/-^* mice showed no difference in motor coordination and strength compared to *Matr3^S85C/S85C^ Clec7a^+/+^* mice as assessed by a variety of motor function tests, including the accelerating rotarod test, inverted grid test, righting reflex, and gait analysis (Figures 2A-2E). Additionally, *Matr3^S85C/S85C^ Clec7a^-/-^* mice showed no difference in weight, body condition score, or hindlimb clasping score compared to *Matr3^S85C/S85C^ Clec7a^+/+^* mice (Figure S3). We also assessed the age at which these mice reached humane endpoint and found no significant difference in survival of *Matr3^S85C/S85C^ Clec7a^-/-^* when compared to *Matr3^S85C/S85C^ Clec7a^+/+^* mice (Figure 2F). These results suggest that Dectin-1 does not play a disease-modifying role in ALS.

**Figure 2.**
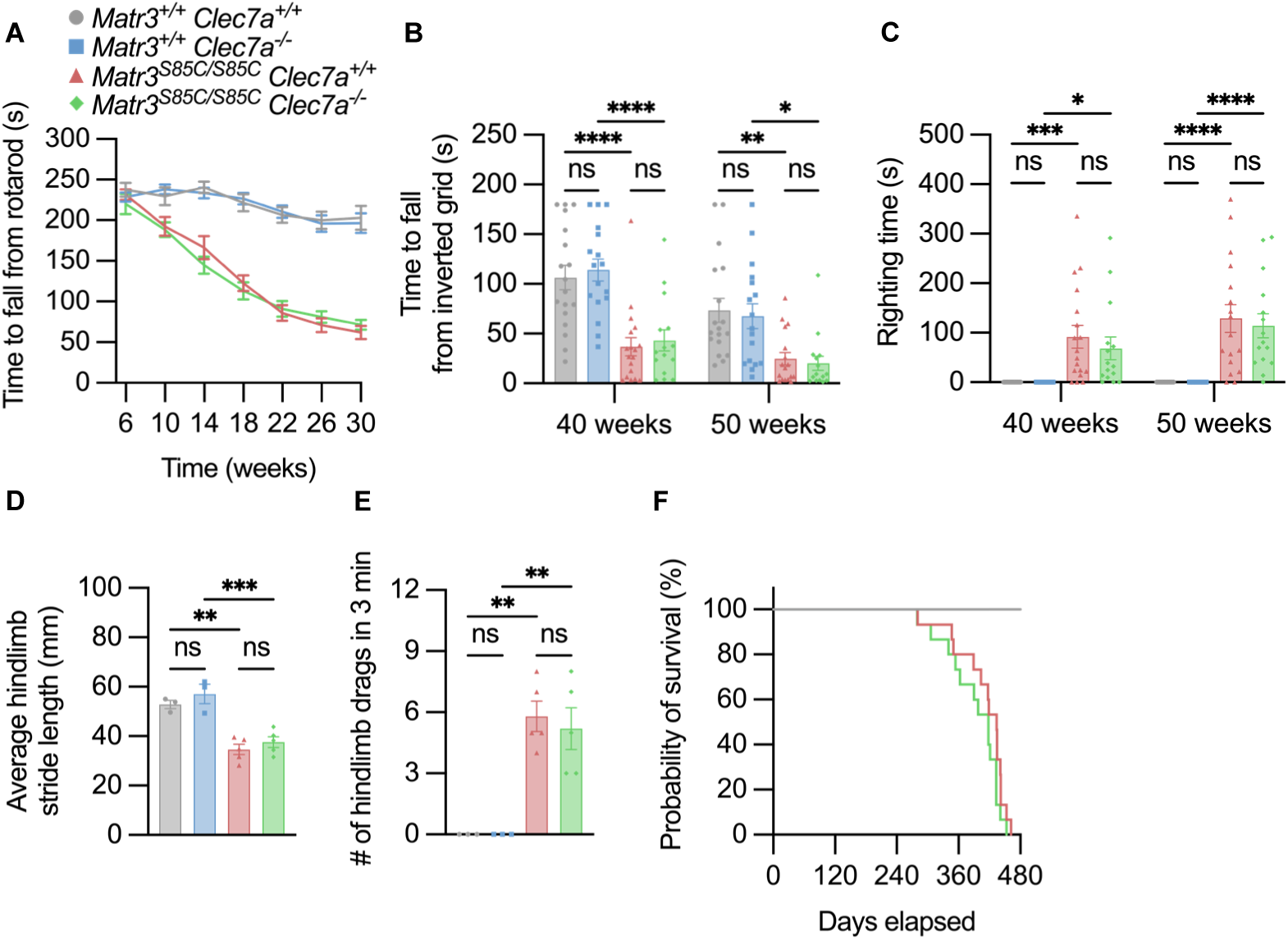
Knockout of Dectin-1 does not affect motor function or survival in *Matr3^S85C/S85C^* mice. (A) Motor coordination and balance was assessed using the Rotarod test from 6 to 30 weeks of age with 4-week intervals (*Matr3^+/+^ Clec7a^+/+^*, *n* = 19; *Matr3^+/+^ Clec7a^-/-^*, *n* = 20; *Matr3^S85C/S85C^ Clec7a^+/+^*, *n* = 18; *Matr3^S85C/S85C^ Clec7a^-/-^*, *n* = 18). (B) Muscle strength was tested using the inverted grid test at 40 weeks of age (*Matr3^+/+^ Clec7a^+/+^*, *n* = 18; *Matr3^+/+^ Clec7a^-/-^*, *n* = 17; *Matr3^S85C/S85C^ Clec7a^+/+^*, *n* = 18; *Matr3^S85C/S85C^ Clec7a^-/-^*, *n* = 15) and 50 weeks of age (*Matr3^+/+^ Clec7a^+/+^*, *n* = 18; *Matr3^+/+^ Clec7a^-/-^*, *n* = 17; *Matr3^S85C/S85C^ Clec7a^+/+^*, *n* = 17; *Matr3^S85C/S85C^ Clec7a^-/-^*, *n* = 15). Bar heights depict mean ± SEM, with each dot representing a single animal. Significance was determined by an ordinary two-way ANOVA corrected by the Tukey method. (C) Delayed righting reflex was measured at 40 weeks of age (*Matr3^+/+^ Clec7a^+/+^*, *n* = 18; *Matr3^+/+^ Clec7a^-/-^*, *n* = 17; *Matr3^S85C/S85C^ Clec7a^+/+^*, *n* = 18; *Matr3^S85C/S85C^ Clec7a^-/-^*, *n* = 15) and 50 weeks of age (*Matr3^+/+^ Clec7a^+/+^*, *n* = 18; *Matr3^+/+^ Clec7a^-/-^*, *n* = 17; *Matr3^S85C/S85C^ Clec7a^+/+^*, *n* = 17; *Matr3^S85C/S85C^ Clec7a^-/-^*, *n* = 15) and represents the time an animal takes to right itself after being flipped on each side. Bar heights depict mean ± SEM, with each dot representing a single animal. Significance was determined by an ordinary two-way ANOVA corrected by the Tukey method. (D) The average hindlimb stride length was measured for five consecutive steps from two replicates at approximately 60 weeks of age (*Matr3^+/+^ Clec7a^+/+^*, *n* = 3; *Matr3^+/+^ Clec7a^-/-^*, *n* = 3; *Matr3^S85C/S85C^ Clec7a^+/+^*, *n* = 5; *Matr3^S85C/S85C^ Clec7a^-/-^*, *n* = 5). Bar heights depict mean ± SEM, with each dot representing a single animal. Significance was determined by an ordinary one-way ANOVA corrected by the Šídák method. (E) The number of times a mouse drags their hindlimbs in 3 min at approximately 60 weeks of age (*Matr3^+/+^ Clec7a^+/+^*, *n* = 3; *Matr3^+/+^ Clec7a^-/-^*, *n* = 3; *Matr3^S85C/S85C^ Clec7a^+/+^*, *n* = 5; *Matr3^S85C/S85C^ Clec7a^-/-^*, *n* = 5). Bar heights depict mean ± SEM, with each dot representing a single animal. Significance was determined by an ordinary one-way ANOVA corrected by the Šídák method. (F) Survival curve (*Matr3^+/+^ Clec7a^+/+^*, *n* = 19; *Matr3^+/+^ Clec7a^-/-^*, *n* = 20; *Matr3^S85C/S85C^ Clec7a^+/+^*, *n* = 15; *Matr3^S85C/S85C^ Clec7a^-/-^*, *n* = 15). Significance was determined by a logrank test.

### Knockout of Dectin-1 does not affect neurodegeneration or microglial responses in *Matr3^S85C/S85C^* mice

We have previously shown that *Matr3^S85C/S85C^* mice exhibit Purkinje cell loss, defects in lower motor neurons, and increased microglial responses^29,30^. Therefore, we performed immunohistochemistry to assess whether knockout of Dectin-1 affects these neuropathological phenotypes. In line with our previous findings^29^, *Matr3^S85C/S85C^ Clec7a^+/+^* mice exhibit significantly fewer Purkinje cells in the cerebellum and an increased percentage of partial denervation of neuromuscular junctions in the tibialis anterior muscle compared to *Matr3^+/+^ Clec7a^+/+^* mice at 60 weeks of age (disease end stage) (Figure 3). However, we did not observe any differences in the number of Purkinje cells or percentage of partial denervation in neuromuscular junctions between *Matr3^S85C/S85C^ Clec7a^+/+^* mice and *Matr3^S85C/S85C^ Clec7a^-/-^* mice (Figure 3), demonstrating that loss of Dectin-1 does not impact neuropathology of MATR3 mutant mice. To determine whether loss of Dectin-1 alters the number of DAM in *Matr3^S85C/S85C^* mice, we immunostained the cerebellum at the advanced disease stage (30 weeks of age) using IBA1 as a pan-microglial marker and CD11c as a DAM marker. There was a significant increase in the number of CD11c^+^ cells in the *Matr3^S85C/S85C^ Clec7a^+/+^*cerebellum compared to *Matr3^+/+^ Clec7a^+/+^*, which was accompanied by an increase in IBA1^+^ cell density (Figure 4). However, there were no overt differences in the numbers of CD11c^+^ or IBA1^+^ cells in *Matr3^S85C/S85C^ Clec7a^-/-^* mice compared to *Matr3^S85C/S85C^ Clec7a^+/+^* mice. Taken together, these results suggest that the loss of Dectin-1 does not modify neurodegeneration or microglial responses in ALS.

**Figure 3.**
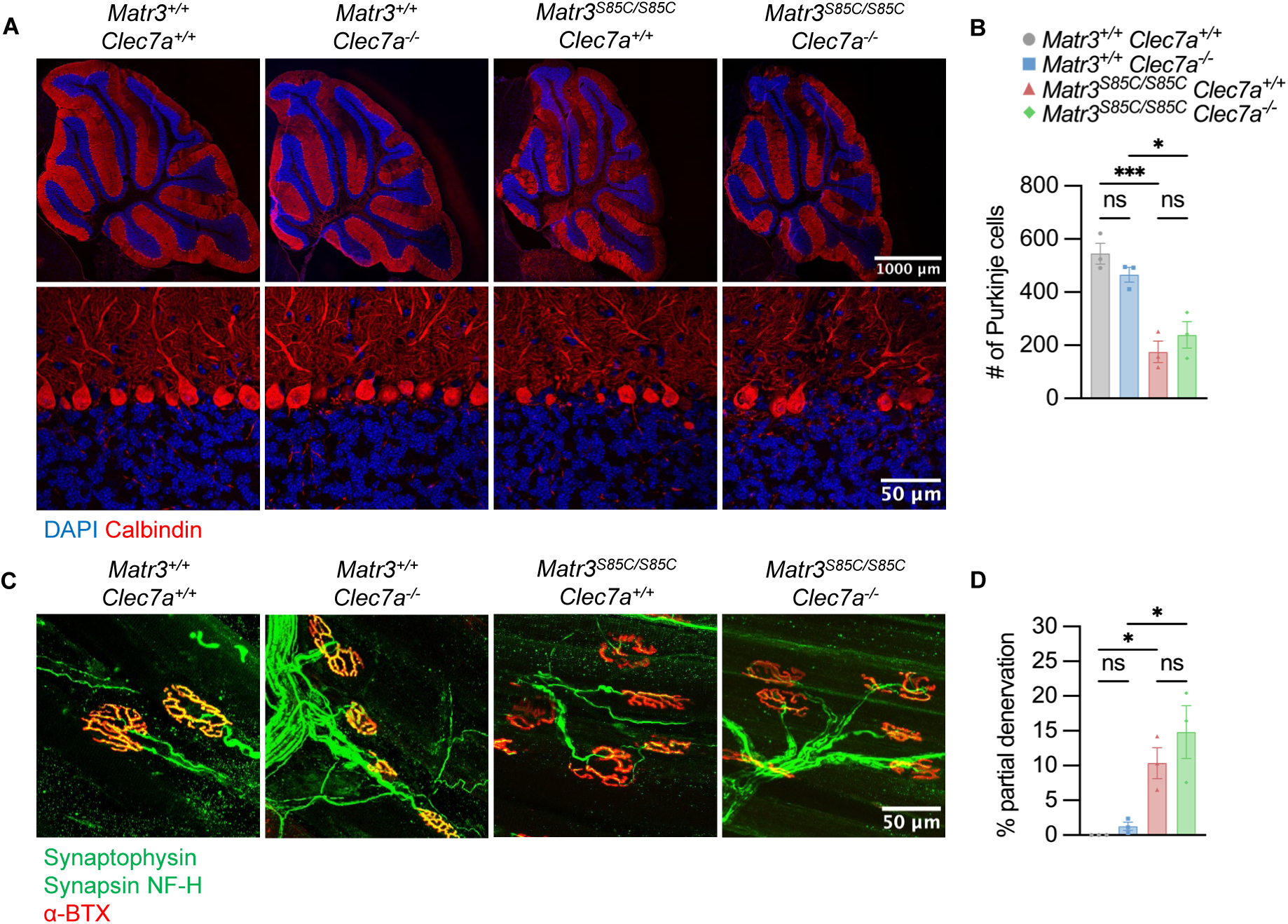
Knockout of Dectin-1 does not affect neurodegeneration in *Matr3^S85C/S85C^* mice. (A) Representative images of the whole cerebellum (upper row) and magnified images of Calbindin^+^ Purkinje cells (bottom row) of *Matr3^+/+^ Clec7a^+/+^*, *Matr3^+/+^ Clec7a^-/-^*, *Matr3^S85C/S85C^ Clec7a^+/+^*, and *Matr3^S85C/S85C^ Clec7a^-/-^* mice at 60 weeks of age. (B) Quantification of the total number of Calbindin^+^ Purkinje cells in a cerebellar section from panel A (*Matr3^+/+^ Clec7a^+/+^*, *n* = 3; *Matr3^+/+^ Clec7a^-/-^*, *n* = 3; *Matr3^S85C/S85C^ Clec7a^+/+^*, *n* = 3; *Matr3^S85C/S85C^ Clec7a^-/-^*, *n* = 3). Bar heights depict mean ± SEM with each dot representing a single animal. Significance was determined by an ordinary one-way ANOVA corrected by the Šídák method. (C) Representative images of neuromuscular junctions in endpoint tibialis anterior showing synaptophysin and synapsin staining (green) of presynaptic terminals, neurofilament H staining (green) of presynaptic axons, and α-bungarotoxin staining (red) of post-synaptic terminals at 60 weeks of age. (D) Quantification of the percentage of neuromuscular junctions in the tibialis anterior muscle exhibiting partial denervation (*Matr3^+/+^ Clec7a^+/+^*, *n* = 3; *Matr3^+/+^ Clec7a^-/-^*, *n* = 3; *Matr3^S85C/S85C^ Clec7a^+/+^*, *n* = 3; *Matr3^S85C/S85C^ Clec7a^-/-^*, *n* = 3). Bar heights depict mean ± SEM with each dot representing a single animal. Significance was determined by an ordinary one-way ANOVA corrected by the Šídák method.

**Figure 4.**
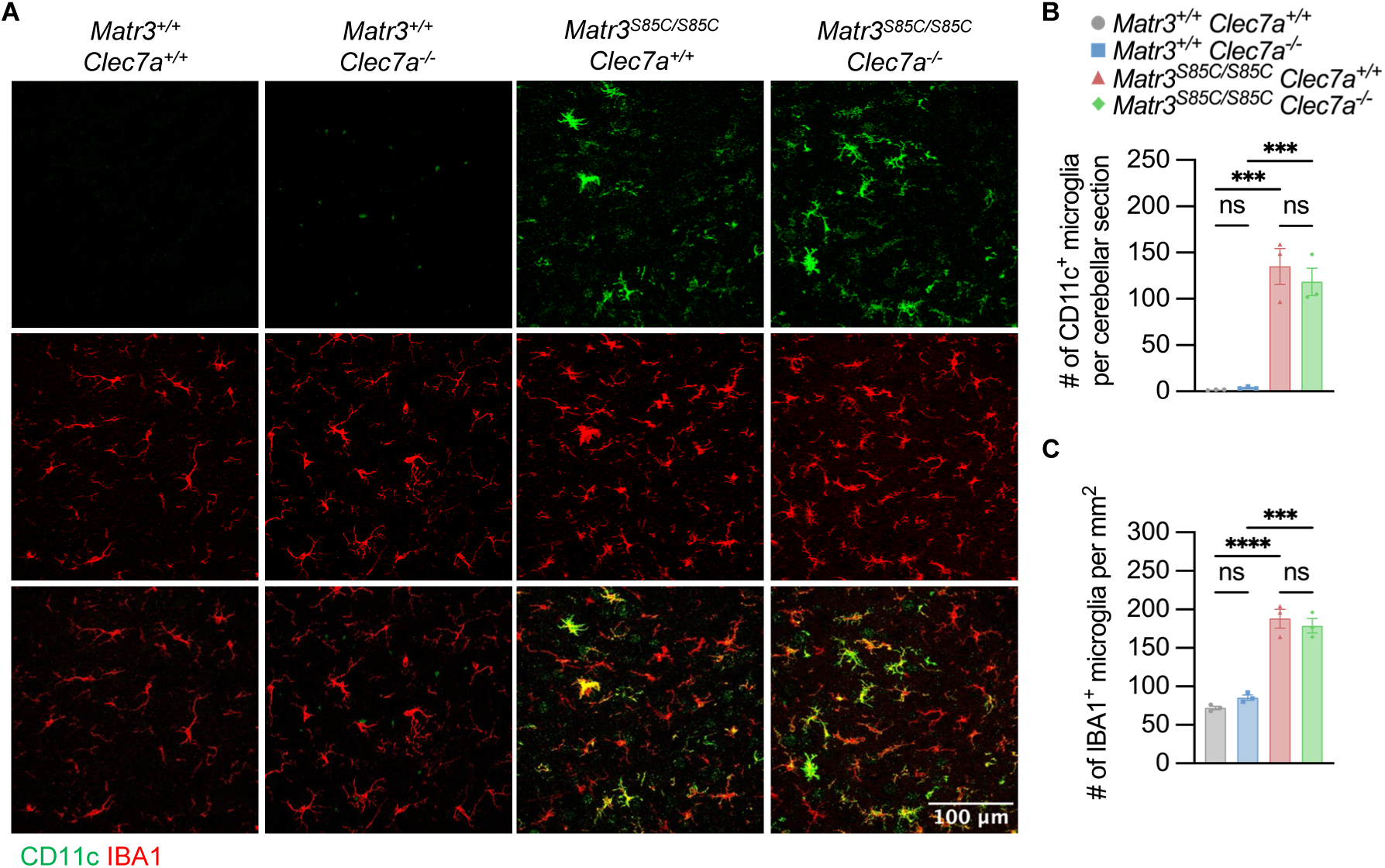
Knockout of Dectin-1 does not affect microglial responses in *Matr3^S85C/S85C^* mice. (A) Representative images of the deep cerebellar nuclei showing CD11c (green) and IBA1 (red) staining for *Matr3^+/+^ Clec7a^+/+^*, *Matr3^+/+^ Clec7a^-/-^*, *Matr3^S85C/S85C^ Clec7a^+/+^*, and *Matr3^S85C/S85C^ Clec7a^-/-^* mice at 30 weeks. (B) Quantification of the total number of CD11c^+^ microglia per cerebellar section and (C) density of IBA1^+^ microglia in the deep cerebellar nuclei (*Matr3^+/+^ Clec7a^+/+^*, *n* = 3; *Matr3^+/+^ Clec7a^-/-^*, *n* = 3; *Matr3^S85C/S85C^ Clec7a^+/+^*, *n* = 3; *Matr3^S85C/S85C^ Clec7a^-/-^*, *n* = 3). Bar heights depict mean ± SEM with each dot representing a single animal. Significance was determined by an ordinary one-way ANOVA corrected by the Šídák method.

## Discussion

DAM play important roles in the neurodegenerative disease process^3,32,33^. In particular, there has been support for a neuroprotective role of DAM through phagocytosis of by-products that accumulate during neurodegeneration, including apoptotic neurons, myelin debris, and toxic proteins, including amyloid beta (Aβ) or pathological TDP-43^5–8^. Although certain DAM-associated proteins, such as TREM2 and SYK, have been extensively studied in various neurodegenerative disease models, the role of other DAM-associated proteins, such as Dectin-1, are less understood. Here, we sought to investigate the role of Dectin-1 on ALS onset and progression. We found that *Clec7a* is upregulated in the cerebellum and spinal cord, both regions that are affected in *Matr3^S85C/S85C^* mice, at early and advanced disease stages. However, we found no change in the expression level of *Clec7a* mRNA in other brain regions or peripheral tissues. Additionally, we show that high numbers of Dectin-1^+^ microglia appear with the onset of motor deficits and their numbers increase with disease severity. These findings are in line with many previous reports showing that DAM appear in the presence of neurodegenerative cues and that its numbers correlate with disease progression^3^.

Genetic and pharmacological targeting of Dectin-1 modifies many different neurological conditions, including AD^9,18,22^, MS^17^, PD^23^, stroke^24,25^, optic nerve injury^19–21^, and axonal injury^34^. Several of these studies have used a Dectin-1 knockout model, which harbours a deletion of exons 1-3, corresponding to the cytoplasmic tail, transmembrane region, and stalk^31^. Knockout of Dectin-1 has been shown to improve neuroinflammation, neurodegeneration, and cognitive function in AD mice^22^. Furthermore, knockout of Dectin-1 rescued the dieback of corticospinal tract axons in a mouse model of spinal cord injury^34^. Conversely, deletion of Dectin-1 worsened clinical score in MS mice^17^ and blocked the pro-regenerative effects of zymosan and curdlan in a mouse model of optic nerve crush injury^20^. These findings highlight that Dectin-1 knockout is sufficient to modify disease phenotypes in mouse models of neurodegeneration and CNS injury, while showing conflicting roles depending on the disease. To investigate the role of Dectin-1 in ALS, we have crossed this same Dectin-1 knockout mouse line with *Matr3^S85C/S85C^* mice. Unexpectedly, we found that the knockout of Dectin-1 did not significantly affect motor deficits, neurodegeneration, and survival in *Matr3^S85C/S85C^*mice. Furthermore, microglial responses in *Matr3^S85C/S85C^* mice, as measured by the number of CD11c^+^ and IBA1^+^ cells, was not affected upon loss of Dectin-1, which may explain the lack of modified behavioral and neuropathological phenotypes. Our findings suggest that loss of Dectin-1 has no observable effect on the disease onset and progression in MATR3 S85C-associated ALS in mice.

One possible explanation for the lack of a disease-modifying effect resulting from the Dectin-1 knockout could be genetic redundancy in DAM signaling in the context of ALS. In two recent studies, injections of a Dectin-1 agonist were shown to induce a neuroprotective microglial response by increasing phagocytosis of Aβ and increasing the number of DAM^9,18^. Interestingly, these neuroprotective effects were shown to be SYK-dependent. These findings highlight that SYK is an important neuroprotective downstream signaling molecule for microglial Dectin-1. However, TREM2 has also been shown to regulate microglial transition to DAM through an SYK-dependent manner^9,18^. Furthermore, there exist several other receptors that could potentially leverage SYK signaling and modify neurodegeneration, including CD22, CD33, CR3, CLEC16A, CLECL1, CLEC2D, FCRL2, and FCRL3^18,35–40^. Our data suggests a possible genetic redundancy of Dectin-1 in neuroprotective DAM functions, at least in ALS, given that the Dectin-1 knockout did not affect disease progression or the number of CD11c^+^ DAM in *Matr3^S85C/S85C^* mice. Future studies should investigate whether increasing Dectin-1-SYK signaling in ALS models could increase neuroprotective DAM activity.

Together, our findings implicate that Dectin-1 does not modify ALS onset or progression but could serve as a useful biomarker. This study may provide insights into the growing body of knowledge on the role of Dectin-1 in neurodegenerative diseases. Our findings highlight the need for more research on the role of Dectin-1 to understand its various (beneficial, detrimental, or neutral) effects upon loss of Dectin-1 or activation of Dectin-1-SYK signaling in various neurodegenerative diseases.

### Limitations of the study

To understand the role of DAM in ALS, it would be worthwhile to assess the effects of stimulating Dectin-1 signaling, such as through a pharmacological agonist, at the onset and during the progression of ALS. It is possible that Dectin-1 knockout does not affect DAM activity due to genetic redundancy, whereas the treatment with a Dectin-1 agonist can boost neuroprotective DAM responses, as shown in AD mice^9,18^. Furthermore, this approach could also potentially modify peripheral immune cells important for disease progression. Studies investigating the role of Dectin-1 in mouse models of MS and optic nerve injury have shown that injections with Dectin-1 agonists can induce an anti-inflammatory or pro-regenerative effect on CNS-infiltrating myeloid cells^17,21^. Therefore, it would be interesting to study whether injection of a Dectin-1 agonist rescues the phenotypes of *Matr3^S85C/S85C^* mice and how DAM, peripheral immune cells and CNS infiltrating cells are playing beneficial roles to the disease. In addition, crossing a *Trem2* knockout or a microglial-specific *Syk* knockout with *Matr3^S85C/S85C^* mice would allow the assessment of consequences due to the ablation of DAM signaling in an ALS mouse model.

## Acknowledgements

We thank the members of the Park laboratory and Dr. Monica Justice laboratory for invaluable discussions on this project. We appreciate the support from the Centre for Phenogenomics (TCP) for mouse care and maintenance, histology, and behavioral phenotyping. We also thank SickKids imaging facility for the use of the microscopes. We thank the lab of Dr. Ronald Cohn for the use of their cryostat. This study was supported by Canadian Institutes of Health Research (CIHR), Brain Canada Foundation Future Leaders Program, Frick Foundation and Canada Research Chairs Program funds to J. P. J. L. was supported by the CIHR Canada Graduate Scholarship, University of Toronto Fellowship and SickKids Restracomp Scholarship. J. R. S. was supported by a Catalyzing the Talent Pipeline Scholarship from the David Dime Family Catalyst Initiative in Molecular Genetics at the University of Toronto. This paper was partly adapted from Justin You’s Master of Science thesis conferred from the University of Toronto.

## Author Contributions

J. Y., M. W. S. and J. P. conceptualized the project. J. Y. and J. P. designed the experiments. J. Y., K. M., K. C., J. L., A. S., J. R. S., and M. M. M. Y. performed the experiments and obtained the data. J. Y. analyzed and interpreted the results. M. W. S. and J. P. supervised the research. J. Y. and J. P. wrote the manuscript. All authors edited, read, and approved the manuscript.

## Declaration of interests

The authors declare no competing interests.

## Methods

### Animals

All mouse procedures were performed under the approval of the Animal Care Committee (ACC) and housed at The Centre for Phenogenomics (TCP), accredited by the Association for Assessment and Accreditation of Laboratory Animal Care International. Mice were kept on a 12 h light/12 h dark cycle at 20-22°C and 43% humidity, with ad libitum access to food and water. *Matr3^S85C/S85C^*mice were developed as previously reported^29^. *Clec7a^-/-^*mice were previously generated by Dr. Gordon Brown^31^ and obtained from Jackson laboratory (012337).

### Monitoring of body weight and health decline

Starting at 4 weeks of age, the body weight of all mice was measured bi-weekly. Concurrently, animals were scored for body condition, mobility, ambulation, activity level, attitude, behaviour, appearance, hydration, respiration, and ataxia as previously described with slight modifications^29^. Humane endpoint was used instead of natural death to ensure animal welfare. If the combined score exceeds 60, the mice are considered to have reached humane endpoint and are euthanized. Mice are automatically considered to have reached humane endpoint if they cannot right themselves within 2 min. Figure S3C shows the body condition score that was defined as a score of 0 when mice are normal and well-conditioned, 8 if they are under-conditioned with their vertebrae column evident and pelvis palpable, and 14 if the mice are emaciated with a prominent skeletal structure and distinctly segmented vertebrae. Figure S3D shows the hindlimb clasping score that was defined as a score of 0 if both limbs are splayed outwards, 1 if one hindlimb is partially retracted towards the abdomen, 2 if both hindlimbs are partially retracted towards the abdomen, and 3 if both hindlimbs are completely retracted and touching the abdomen.

### Rotarod test

The rotarod test was performed to measure motor coordination, balance, and motor learning ability as previously described^29^. In brief, mice were placed on the rotarod apparatus (Ugo Basile, Stoelting) that accelerated linearly from 5 rpm to 40 rpm in 5 min. The average latency to fall of 3 trials was recorded every 4 weeks from 6 to 30 weeks of age.

### Inverted grid test

The inverted grid test was performed to measure muscle strength as previously described^29^. In brief, mice were placed on a metal grid held approximately 30 cm above the bench. After 3 s, the grid was inverted so that the mice were hanging upside down. The average latency to fall (not exceeding 3 minutes) of 3 trials was recorded for mice at 40 and 50 weeks of age.

### Delayed righting test

The delayed righting test was performed to measure the righting reflex as previously described^29^. In brief, mice were placed onto their side in their home cage with cage mates present. The latency for the mice to right themselves for each side was measured and summed at 40 and 50 weeks of age.

### Footprint test

The footprint test was performed to measure gait impairment as previously described with minor modifications^29^. In brief, the hind paws were painted with non-toxic dye and the mice were allowed to freely walk along a paper-covered runway. The distance for each footprint was measured to obtain an average for five consecutive steps. Two replicates were obtained per mouse.

### Hindlimb dragging test

The hindlimb dragging test was performed to measure gait impairment as previously described^29^. In brief, the number of times that freely moving mice exhibited hindlimb dragging was recorded over 3 min.

### RNA extraction, cDNA synthesis, and RT-PCR

The cerebellum, spinal cord, forebrain/brainstem, spleen, and tibialis anterior muscle were flash frozen and stored at -80°C. The samples and 1 mL of TRIzol reagent (Invitrogen, 15596018) were added to tubes filled with 1.4 mm Zirconium beads (OPS Diagnostics, PFAW 1400-100-19) and homogenized using the MagNA Lyser (Roche) at 7,000 rpm for 20 s. The solution was transferred into a new 1.5 mL tube and the samples were incubated at RT for 3 min. 200 μL of chloroform (Sigma-Aldrich, 67-66-3) was added to each tube and vortexed. The samples were incubated at RT for 10 min and then centrifuged at 14,000 rpm at 4°C for 20 min. The upper clear layer was transferred into new tubes containing 500 μL of isopropanol and was inverted to mix. The samples were incubated at RT for 10 min and then centrifuged at 14,000 rpm at 4°C for 20 min. The supernatant was discarded, and the pellet was cleaned with 500 μL of 75% ethanol. The samples were centrifuged at 14,000 rpm at 4°C for 5 min. The supernatant was discarded, and the pellet was dried at RT for 10 min. The RNA pellet was resuspended in 50-100 μL of DEPC-treated water (Invitrogen, 4387937) and incubated at 55°C for 15 min. RNA was kept at - 80°C for long-term storage.

RNA concentrations were measured using the NanoDrop 2000/2000C Spectrophotometer (Thermo Scientific, ND-2000). 2 μg of RNA was combined with nuclease-free water to a total volume of 9 μL, followed by 1 μL of 50 μM random hexamer (Invitrogen, N8080127). The samples were incubated at 70°C for 5 min and then chilled on ice for 5 min. 15 μL of master mix was added to each sample to a final concentration of 0.8X First Strand Buffer (Invitrogen, 28025013), 0.4 mM dNTP mix (New England Biolabs, N0447L), 0.4 U/μL RNaseOUT (Invitrogen, 10777019), 8 U/μL MLV RT (Invitrogen, 28025013) and 1 mM DTT (Invitrogen, 28025013). The samples were incubated at 37°C for 1 h for cDNA generation. The samples were incubated at 95°C for 10 min for heat inactivation of the reverse transcriptase. cDNA was stored at -20°C.

Reverse transcription PCR (RT-PCR) to measure *Clec7a* and *Gapdh* transcript levels was performed using KOD Hot Start DNA Polymerase (VWR, CA80511-386). The following primers were used for RT-PCR: *Clec7a* exons 2-3 (Fwd 5’-CTTCACCTTGGAGGCCCATT -3’; Rev 5’- GAGGGAGCCACCTTCTCATC-3’), *Clec7a* exons 4-5 (Fwd 5’- CTTGCCTTCCTAATTGGATC-3’; Rev 5’-GCATTAATACGGTGAGACGATG-3’), and *Gapdh* (Fwd 5′-ACTCCACTCACGGCAAATTC-3′; Rev 5′-CCTTCCACAATGCCAAAGTT- 3′).

### Immunohistochemistry

Mice were deeply anesthetized with isoflurane, followed by transcardial perfusion with 1X phosphate-buffered saline (PBS) and then 4% paraformaldehyde (Sigma-Aldrich, 30525-89-4) diluted in 1X PBS. After perfusion, the brain was dissected and fixed in 4% paraformaldehyde at RT for 48 h, followed by 3 washes in 1X PBS for 15 min each. The brain was cryoprotected in 30% sucrose and each half of the brain was embedded for sagittal sections in Optimal Cutting Temperature (OCT) compound. 40 μm brain sections were washed 3 times with 0.3% Triton X-100 (Sigma-Aldrich) in 1X PBS (PBST) for 5 min each. For Calbindin staining, 20 μm sections were used instead for better antibody penetration. Sections were blocked in 5% normal donkey serum in 0.3% PBST at RT for 1 h. Sections were incubated with primary antibody diluted in blocking buffer at 4°C for 48 h. The following primary antibodies were used: anti-Dectin-1 (rat monoclonal, Invivogen, mabg-mdect, 1:100), anti-IBA1 (goat polyclonal, Abcam, ab5076, 1:500), anti-CD11c (rabbit monoclonal, Cell Signaling Technology, 97585S, 1:100), and anti-calbindin (mouse monoclonal, Swant, 300, 1:300). Sections were washed 3 times with 0.3% PBST and incubated with secondary antibodies diluted in blocking buffer at RT for 3 h. The following secondary antibodies were used: 488 Alexa Fluor donkey anti-rat (Invitrogen, A-21208, 1:500), 555 Alexa Fluor donkey anti-rabbit (Invitrogen, A-31572, 1:500), and 647 Alexa Fluor donkey anti-goat (Invitrogen, A21447, 1:500). Sections were washed in 0.3% PBST and incubated with DAPI (Millipore, D6210, 1:1000) for 15 min. Sections were washed with 0.3% PBST and mounted with ProLong Gold Antifade mountant (Invitrogen, P36930). All slides were imaged using NIS-Elements software on a Nikon A1R confocal microscope (Nikon Instruments). The number of Calbindin^+^, Dectin-1^+^, IBA1^+^ and CD11c^+^ cells were manually counted.

### Photobleaching

Photobleaching was performed prior to immunostaining of microglial markers to remove lipofuscin-like autofluorescence^41^. Photobleaching was performed using a lab-made photobleacher as previously described^41^. After sectioning, floating sections submerged in 1X PBS were placed under a blue LED light bulb for 24 hours to photobleach and then proceeded for immunostaining as described above.

### Whole-mount immunostaining of tibialis anterior (TA) muscle

Following transcardial perfusion, the tibialis anterior (TA) muscle was fixed in 4% paraformaldehyde for 15 minutes, then washed 3 times for 5 minutes each in 1X PBS. TA muscles were teased into thin pieces with the membrane and fat removed. Muscle tissue was incubated in 0.1 M glycine in 1X PBS (pH 3.0) for 30 minutes at RT, rocking gently, to block free aldehydes. Muscle tissue was then permeabilized in 2% Triton X-100 in 1X PBS for 30 minutes at RT, rocking gently. Tissues were blocked in blocking buffer comprised of 4% Bovine Serum Albumin (Roche, 10735087001) and 1% Triton X-100 in 1X PBS for 30 minutes at RT, rocking gently. Tissues were then incubated with primary antibody diluted in blocking buffer overnight at 4°C. The following primary antibodies were used: synaptophysin antibody (rabbit polyclonal, Synaptic Systems 101 002, 1:200), synapsin I antibody (rabbit polyclonal, ab64581, 1:200), and neurofilament H antibody (chicken polyclonal, ab4680, 1:200). Tissues were washed with 1X PBS for 3 times, 30 minutes each. The tissues were incubated in secondary antibodies, α-bungarotoxin (Alexa Fluor 555 conjugate, Invitrogen B35451, 1:500) and DAPI (Millipore D6210, 1:1000) for 2 hours at RT, rocking gently. The following secondary antibodies were used: 488 Alexa Fluor donkey anti-rabbit (Invitrogen, A-21206, 1:250) and 488 Alexa Fluor goat anti-chicken (Invitrogen, A-11039, 1:250). Tissues were washed with 1X PBS for 3 times, 10 minutes each, then mounted on concavity slides using ProLong Gold Antifade mountant (Invitrogen, P36930). All slides were imaged using SP8 Leica confocal microscope (Leica Microsystems).

### Neuromuscular junction quantification

Teased TA muscle was stained and imaged. About 20 to 30 confocal z-series were imaged per animal at 40X magnification, 0.75 zoom, and a step size of 3 µm. Over 100 neuromuscular junctions with connections to presynaptic axons were imaged per animal. The number of neuromuscular junctions with presynaptic axonal swelling (blebbing) was manually counted. Of the neuromuscular junctions that were visibly connected to presynaptic axons, endplates showing approximately less than 80% but greater than 1% overlap with presynaptic terminal staining were classified as partially denervated and were manually counted.

### Statistical analysis

Statistical significance was determined using an ordinary one-way ANOVA corrected for multiple comparisons using the Šídák method, ordinary two-way ANOVA corrected using the Šídák method, ordinary two-way ANOVA corrected using the Tukey method, or logrank test, as indicated in the figure legends. All statistical tests were performed using GraphPad Prism 10.1.0: \**P* < 0.05, \*\**P* < 0.01, \*\*\**P* < 0.001, \*\*\*\**P* < 0.0001, ns = not significant.

## Notes

### Competing Interest Statement

The authors have declared no competing interest.

## References

1. Hammond, T. R. et al. Single-Cell RNA Sequencing of Microglia throughout the Mouse Lifespan and in the Injured Brain Reveals Complex Cell-State Changes. Immunity 50, 253–271.e6 (2019).

2. Li, Q. et al. Developmental Heterogeneity of Microglia and Brain Myeloid Cells Revealed by Deep Single-Cell RNA Sequencing. Neuron 101, 207–223.e10 (2019).

3. Deczkowska, A. et al. Disease-Associated Microglia: A Universal Immune Sensor of Neurodegeneration. Cell 173, 1073–1081 (2018).

4. Paolicelli, R. C. et al. Microglia states and nomenclature: A field at its crossroads. Neuron 110, 3458–3483 (2022).

5. Keren-Shaul, H. et al. A Unique Microglia Type Associated with Restricting Development of Alzheimer’s Disease. Cell 169, 1276–1290.e17 (2017).

6. Krasemann, S. et al. The TREM2-APOE Pathway Drives the Transcriptional Phenotype of Dysfunctional Microglia in Neurodegenerative Diseases. Immunity 47, 566–581.e9 (2017).

7. Xie, M. et al. TREM2 interacts with TDP-43 and mediates microglial neuroprotection against TDP-43-related neurodegeneration. Nat. Neurosci. 25, 26–38 (2022).

8. Nugent, A. A. et al. TREM2 Regulates Microglial Cholesterol Metabolism upon Chronic Phagocytic Challenge. Neuron 105, 837–854.e9 (2020).

9. Wang, S. et al. TREM2 drives microglia response to amyloid-β via SYK-dependent and - independent pathways. Cell 185, 4153–4169.e19 (2022).

10. Safaiyan, S. et al. White matter aging drives microglial diversity. Neuron 109, 1100–1117.e10 (2021).

11. Wang, Y. et al. TREM2 Lipid Sensing Sustains the Microglial Response in an Alzheimer’s Disease Model. Cell 160, 1061–1071 (2015).

12. Lee, S.-H. et al. Trem2 restrains the enhancement of tau accumulation and neurodegeneration by β-amyloid pathology. Neuron 109, 1283–1301.e6 (2021).

13. Lee, C. Y. D. et al. Elevated TREM2 Gene Dosage Reprograms Microglia Responsivity and Ameliorates Pathological Phenotypes in Alzheimer’s Disease Models. Neuron 97, 1032–1048.e5 (2018).

14. Brown, G. D., Willment, J. A. & Whitehead, L. C-type lectins in immunity and homeostasis. Nat. Rev. Immunol. 18, 374–389 (2018).

15. Tone, K., Stappers, M. H. T., Willment, J. A. & Brown, G. D. C-type lectin receptors of the Dectin-1 cluster: Physiological roles and involvement in disease. Eur. J. Immunol. 49, 2127–2133 (2019).

16. Deerhake, M. E. & Shinohara, M. L. Emerging roles of Dectin-1 in noninfectious settings and in the CNS. Trends Immunol. 42, 891–903 (2021).

17. Deerhake, M. E. et al. Dectin-1 limits autoimmune neuroinflammation and promotes myeloid cell-astrocyte crosstalk via Card9-independent expression of Oncostatin M. Immunity 54, 484–498.e8 (2021).

18. Ennerfelt, H. et al. SYK coordinates neuroprotective microglial responses in neurodegenerative disease. Cell 185, 4135–4152.e22 (2022).

19. Yin, Y. et al. Macrophage-derived factors stimulate optic nerve regeneration. J. Neurosci. Off. J. Soc. Neurosci. 23, 2284–2293 (2003).

20. Baldwin, K. T., Carbajal, K. S., Segal, B. M. & Giger, R. J. Neuroinflammation triggered by β-glucan/dectin-1 signaling enables CNS axon regeneration. Proc. Natl. Acad. Sci. U. S. A. 112, 2581–2586 (2015).

21. Sas, A. R. et al. A new neutrophil subset promotes CNS neuron survival and axon regeneration. Nat. Immunol. 21, 1496–1505 (2020).

22. Zhao, X. et al. β-amyloid binds to microglia Dectin-1 to induce inflammatory response in the pathogenesis of Alzheimer’s disease. Int. J. Biol. Sci. 19, 3249–3265 (2023).

23. Chen, X.-Y., Feng, S.-N., Bao, Y., Zhou, Y.-X. & Ba, F. Identification of Clec7a as the therapeutic target of rTMS in alleviating Parkinson’s disease: targeting neuroinflammation. Biochim. Biophys. Acta Mol. Basis Dis. 1869, 166814 (2023).

24. Ye, X.-C. et al. Dectin-1/Syk signaling triggers neuroinflammation after ischemic stroke in mice. J. Neuroinflammation 17, 17 (2020).

25. Fu, X. et al. Inhibition of Dectin-1 Ameliorates Neuroinflammation by Regulating Microglia/Macrophage Phenotype After Intracerebral Hemorrhage in Mice. Transl. Stroke Res. 12, 1018–1034 (2021).

26. Johnson, J. O. et al. Mutations in the Matrin 3 gene cause familial amyotrophic lateral sclerosis. Nat. Neurosci. 17, 664–666 (2014).

27. Zhao, M., Kim, J. R., van Bruggen, R. & Park, J. RNA-Binding Proteins in Amyotrophic Lateral Sclerosis. Mol. Cells 41, 818–829 (2018).

28. Pollini, D. et al. Multilayer and MATR3-dependent regulation of mRNAs maintains pluripotency in human induced pluripotent stem cells. iScience 24, 102197 (2021).

29. Kao, C. S. et al. Selective neuronal degeneration in MATR3 S85C knock-in mouse model of early-stage ALS. Nat. Commun. 11, 5304 (2020).

30. You, J. et al. Selective Loss of MATR3 in Spinal Interneurons, Upper Motor Neurons and Hippocampal CA1 Neurons in a MATR3 S85C Knock-In Mouse Model of Amyotrophic Lateral Sclerosis. Biology 11, 298 (2022).

31. Taylor, P. R. et al. Dectin-1 is required for beta-glucan recognition and control of fungal infection. Nat. Immunol. 8, 31–38 (2007).

32. Hickman, S., Izzy, S., Sen, P., Morsett, L. & El Khoury, J. Microglia in neurodegeneration. Nat. Neurosci. 21, 1359–1369 (2018).

33. You, J., Youssef, M. M. M., Santos, J. R., Lee, J. & Park, J. Microglia and Astrocytes in Amyotrophic Lateral Sclerosis: Disease-Associated States, Pathological Roles, and Therapeutic Potential. Biology 12, 1307 (2023).

34. Gensel, J. C. et al. Toll-Like Receptors and Dectin-1, a C-Type Lectin Receptor, Trigger Divergent Functions in CNS Macrophages. J. Neurosci. Off. J. Soc. Neurosci. 35, 9966–9976 (2015).

35. Clark, E. A. & Giltiay, N. V. CD22: A Regulator of Innate and Adaptive B Cell Responses and Autoimmunity. Front. Immunol. 9, 2235 (2018).

36. Hadas, S., Spira, M., Hanisch, U.-K., Reichert, F. & Rotshenker, S. Complement receptor-3 negatively regulates the phagocytosis of degenerated myelin through tyrosine kinase Syk and cofilin. J. Neuroinflammation 9, 166 (2012).

37. Pluvinage, J. V. et al. CD22 blockade restores homeostatic microglial phagocytosis in ageing brains. Nature 568, 187–192 (2019).

38. Wißfeld, J. et al. Deletion of Alzheimer’s disease-associated CD33 results in an inflammatory human microglia phenotype. Glia 69, 1393–1412 (2021).

39. International Multiple Sclerosis Genetics Consortium. Multiple sclerosis genomic map implicates peripheral immune cells and microglia in susceptibility. Science 365, eaav7188 (2019).

40. International Multiple Sclerosis Genetics Consortium et al. Genetic risk and a primary role for cell-mediated immune mechanisms in multiple sclerosis. Nature 476, 214–219 (2011).

41. Stillman, J. M. et al. Lipofuscin-like autofluorescence within microglia and its impact on studying microglial engulfment. Nat. Commun. 14, 7060 (2023).

