## Supplementary Figures for "Knockout of Dectin-1 does not modify disease onset or progression in a MATR3 S85C knock-in mouse model of ALS"

Figure S1

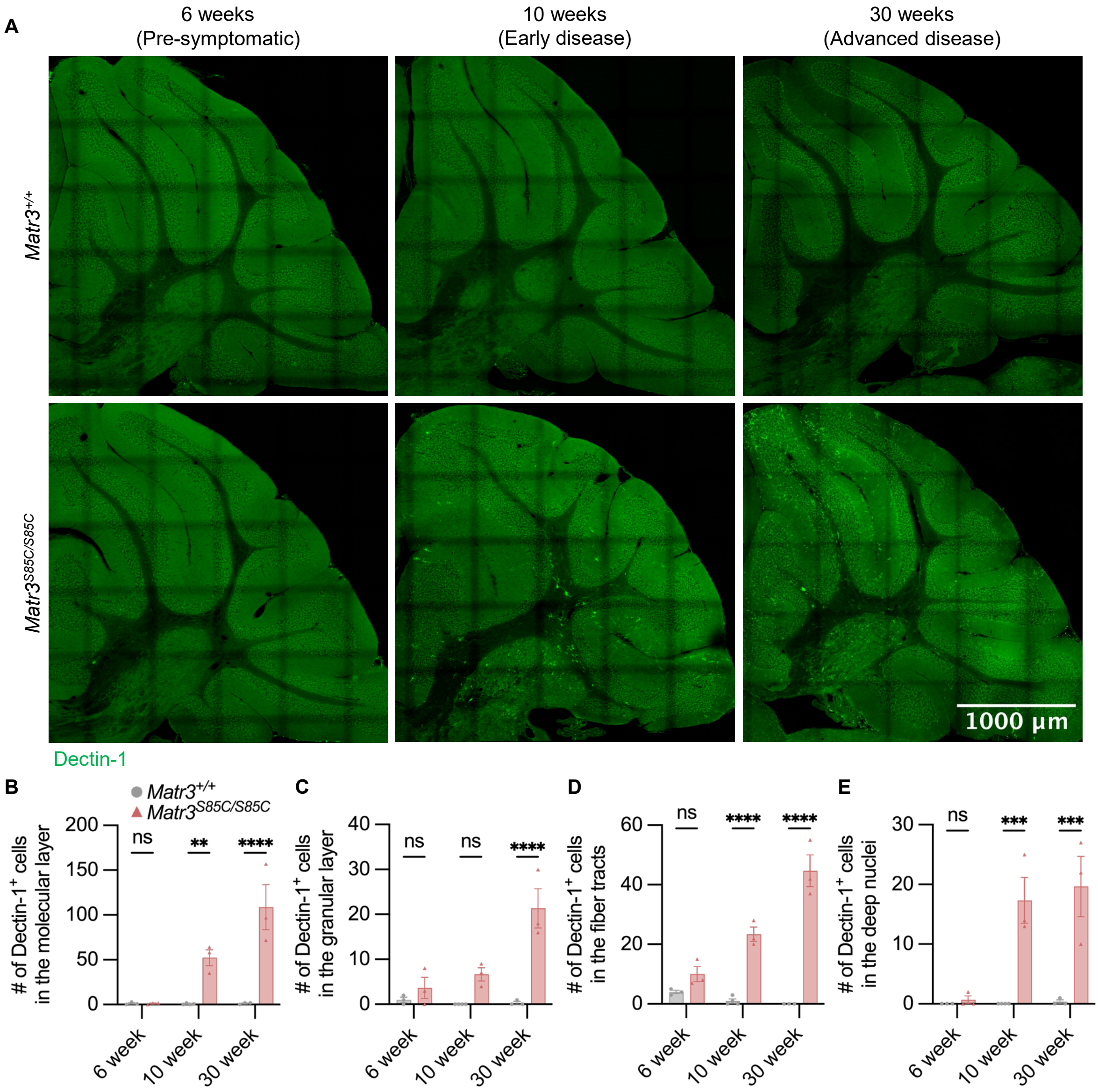

A
6 weeks
(Pre-symptomatic)
10 weeks
(Early disease)
30 weeks
(Advanced disease)
Matr3+/+
Matr3S85C/S85C
1000 µm
Dectin-1
B
Matr3+/+
Matr3S85C/S85C
### of Dectin-1+ cells in the molecular layer
200
150
100
50
0
ns
**
****
6 week
10 week
30 week
C
### of Dectin-1+ cells in the granular layer
40
30
20
10
0
ns
ns
****
6 week
10 week
30 week
D
### of Dectin-1+ cells in the fiber tracts
60
40
20
0
ns
****
****
6 week
10 week
30 week
E
### of Dectin-1+ cells in the deep nuclei
30
20
10
0
ns
***
***
6 week
10 week
30 week

**Figure S1. Increase of Dectin-1$^{+}$ cells throughout the entire cerebellum of *Matr3$^{S85C/S85C}$* mice; Related:to Figure 1".** (A) Representative images of the whole cerebellum showing Dectin-1 staining for *Matr3*$^{+/+}$ and *Matr3*$^{S85C/S85C}$ mice at the indicated ages. (B-E) Quantification of the number of Dectin-1$^{+}$ microglia in the (B) molecular layer, (C) granular layer, (D) fiber tracts, and (E) deep nuclei (*Matr3*$^{+/+}$, $n = 3$; *Matr3*$^{S85C/S85C}$, $n = 3$). Bar heights depict mean ± SEM, with each dot representing a single animal. Significance was determined by an ordinary one-way ANOVA corrected by the Šídák method.

### Figure S2

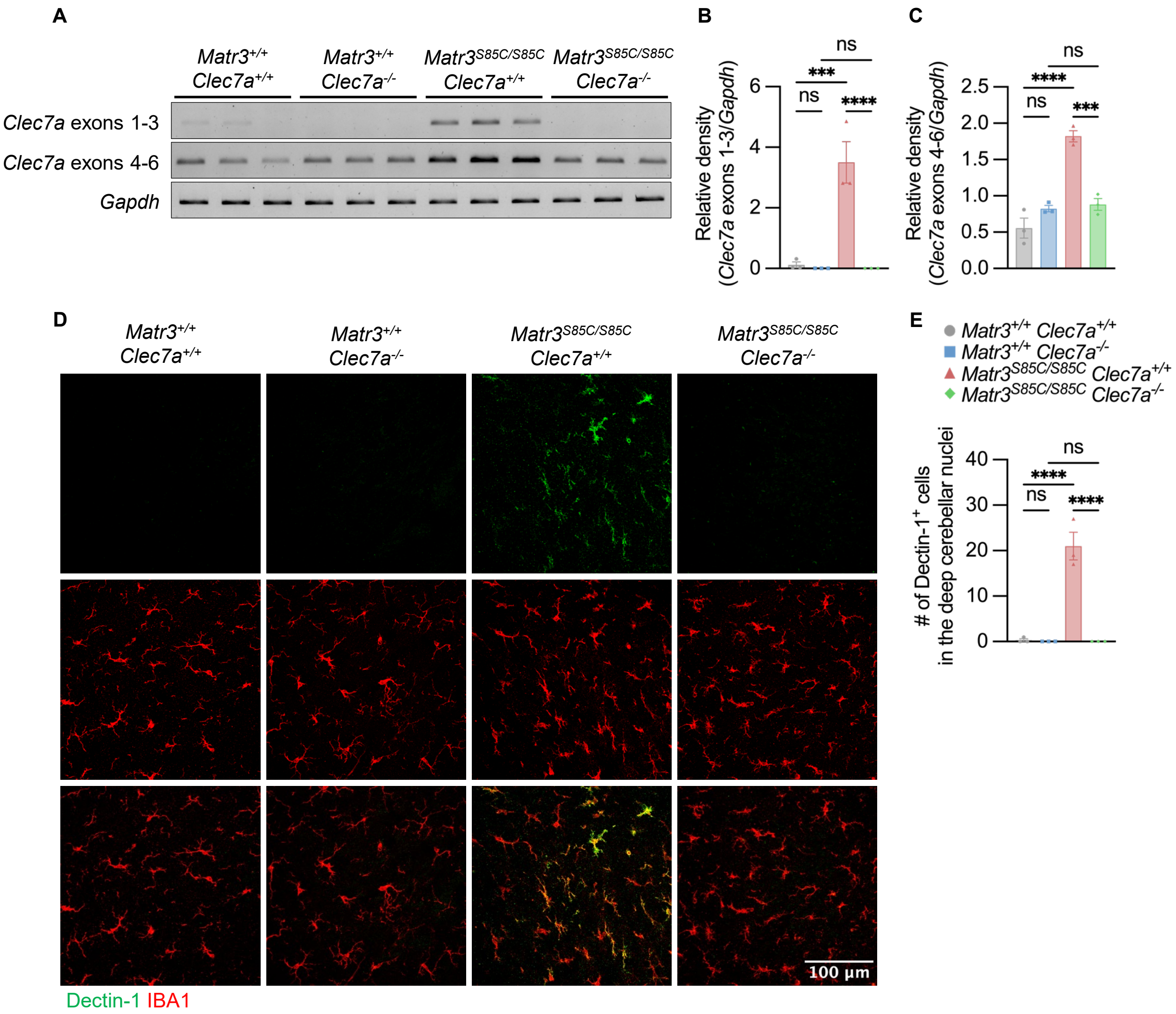

A
Matr3+/+ Clec7a+/+
Matr3+/+ Clec7a-/-
Matr3S85C/S85C Clec7a+/+
Matr3S85C/S85C Clec7a-/-
Clec7a exons 1-3
Clec7a exons 4-6
Gapdh
B
Relative density (Clec7a exons 1-3/Gapdh)
0
2
4
6
ns
***
ns
****
C
Relative density (Clec7a exons 4-6/Gapdh)
0.0
0.5
1.0
1.5
2.0
2.5
ns
****
ns
***
D
Matr3+/+ Clec7a+/+
Matr3+/+ Clec7a-/-
Matr3S85C/S85C Clec7a+/+
Matr3S85C/S85C Clec7a-/-
100 μm
Dectin-1 IBA1
E
Matr3+/+ Clec7a+/+
Matr3+/+ Clec7a-/-
Matr3S85C/S85C Clec7a+/+
Matr3S85C/S85C Clec7a-/-
### of Dectin-1+ cells in the deep cerebellar nuclei
0
10
20
30
40
ns
****
ns
****

**Figure S2. Validation of Dectin-1 knockout in *Matr3*$^{S85C/S85C}$ mice; Related:to Figures 2-4".** (A) Representative images and (B-C) quantification of RT-PCR results showing the mRNA expression levels of *Clec7a* and *Gapdh* from the cerebellum obtained at 30 weeks of age (*Matr3*$^{+/+}$ *Clec7a*$^{+/+}$, *n* = 3; *Matr3*$^{+/+}$ *Clec7a*$^{-/-}$, *n* = 3; *Matr3*$^{S85C/S85C}$ *Clec7a*$^{+/+}$, *n* = 3; *Matr3*$^{S85C/S85C}$ *Clec7a*$^{-/-}$, *n* = 3). *Clec7a* expression is normalized to *Gapdh* expression. Bar heights depict mean ± SEM, with each dot representing a single animal. Significance was determined by an ordinary one-way ANOVA corrected by the Šídák method. (D) Representative images of the deep cerebellar nuclei showing Dectin-1 (green) and IBA1 (red) staining for *Matr3*$^{+/+}$ *Clec7a*$^{+/+}$, *Matr3*$^{+/+}$ *Clec7a*$^{-/-}$, *Matr3*$^{S85C/S85C}$ *Clec7a*$^{+/+}$, and *Matr3*$^{S85C/S85C}$ *Clec7a*$^{-/-}$ mice at 30 weeks. (E) Quantification of the total number of Dectin-1$^{+}$ microglia per deep cerebellar nuclei (*Matr3*$^{+/+}$ *Clec7a*$^{+/+}$, *n* = 3; *Matr3*$^{+/+}$ *Clec7a*$^{-/-}$, *n* = 3; *Matr3*$^{S85C/S85C}$ *Clec7a*$^{+/+}$, *n* = 3; *Matr3*$^{S85C/S85C}$ *Clec7a*$^{-/-}$, *n* = 3). Bar heights depict mean ± SEM, with each dot representing a single animal. Significance was determined by an ordinary one-way ANOVA corrected by the Šídák method.

Figure S3

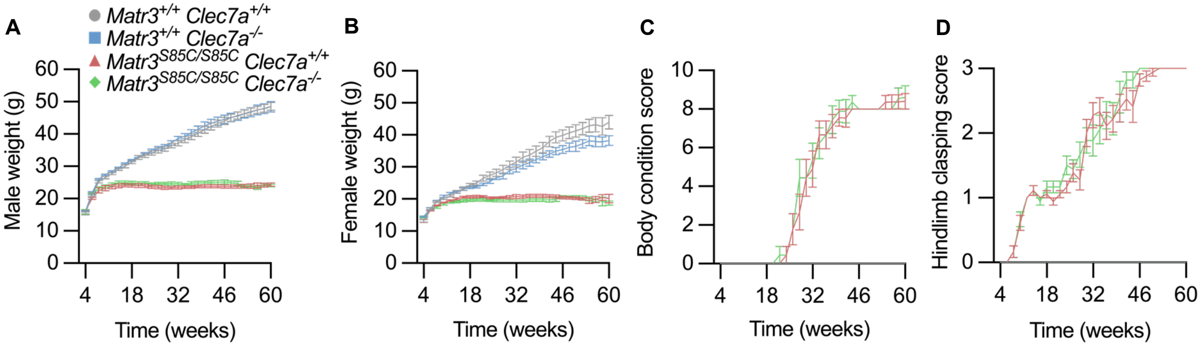


**Figure S3. Knockout of Dectin-1 does not affect body weight, body condition score, or hindlimb clasping score in *Matr3*$^{S85C/S85C}$ mice; Related:to Figure 2".** (A) Graph depicting body weight measured over the lifespan of male mice (*Matr3*$^{+/+}$ *Clec7a*$^{+/+}$, *n* = 10; *Matr3*$^{+/+}$ *Clec7a*$^{-/-}$, *n* = 10; *Matr3*$^{S85C/S85C}$ *Clec7a*$^{+/+}$, *n* = 9; *Matr3*$^{S85C/S85C}$ *Clec7a*$^{-/-}$, *n* = 10) and (B) female mice (*Matr3*$^{+/+}$ *Clec7a*$^{+/+}$, *n* = 9; *Matr3*$^{+/+}$ *Clec7a*$^{-/-}$, *n* = 10; *Matr3*$^{S85C/S85C}$ *Clec7a*$^{+/+}$, *n* = 9; *Matr3*$^{S85C/S85C}$ *Clec7a*$^{-/-}$, *n* = 8). (C) Graph depicting body condition score measured over the lifespan (*Matr3*$^{+/+}$ *Clec7a*$^{+/+}$, *n* = 19; *Matr3*$^{+/+}$ *Clec7a*$^{-/-}$, *n* = 20; *Matr3*$^{S85C/S85C}$ *Clec7a*$^{+/+}$, *n* = 15; *Matr3*$^{S85C/S85C}$ *Clec7a*$^{-/-}$, *n* = 15). (D) Graph depicting hindlimb clasping score measured over the lifespan (*Matr3*$^{+/+}$ *Clec7a*$^{+/+}$, *n* = 19; *Matr3*$^{+/+}$ *Clec7a*$^{-/-}$, *n* = 20; *Matr3*$^{S85C/S85C}$ *Clec7a*$^{+/+}$, *n* = 15; *Matr3*$^{S85C/S85C}$ *Clec7a*$^{-/-}$, *n* = 15). Quantification for panels A-D started at 4 weeks and continued bi-weekly until humane endpoint.
